# EXPLICIT-Kinase: a gene expression predictor for dissecting the functions of the Arabidopsis kinome

**DOI:** 10.1101/2021.09.15.460437

**Authors:** Yuming Peng, Wanzhu Zuo, Yue Qin, Shisong Ma

## Abstract

Protein kinases regulate virtually all cellular processes, but it remains challenging to determine the functions of all protein kinases, collectively called the kinome, in any species. We developed an approach called EXPLICIT-Kinase to predict the functions of the Arabidopsis kinome. Because the activities of many kinases can be regulated transcriptionally, their gene expression patterns provide clues to their functions. A universal gene expression predictor for Arabidopsis was constructed to predict the expression of 30,172 non-kinase genes based on the expression of 994 protein kinase genes. The model reconstituted highly accurate transcriptomes for diverse Arabidopsis samples. It identified the significant kinases as predictor kinases for predicting the expression of Arabidopsis genes and pathways. Strikingly, these predictor kinases were often known regulators of the related pathways, as exemplified by those involved in cytokinesis, tissue development, and stress responses. Comparative analyses have revealed that portions of these predictor kinases, including the novel ones, are shared and conserved between Arabidopsis and maize. The conservation between species provide additional evidence to support the novel predictor kinases as bona fide regulators of the pathways involved. Thus our approach enables the systematic dissection of the functions of the Arabidopsis kinome.

## INTRODUCTION

In eukaryotic species, protein kinases constitute a large and functionally diverse protein superfamily, and all the members within a single species are often referred to collectively as the kinome [1, 2]. Protein kinases modify proteins by adding phosphate groups to them to regulate their activities, localizations, and functions. They are important components of signaling networks and orchestrate almost all cellular processes; examples are cytokinesis, human angiogenesis, plant guard cell movement, and pollen tube growth. Dissecting the functions of the kinome is crucial to understanding the regulatory landscapes of biological systems.

Protein kinases have mostly been studied either individually or within the scope of a small kinase family (e.g. mitogen-activated protein kinases, MAPKs), which falls short of providing a complete coverage of all protein kinases. Systems biology approaches were developed over the past two decades to investigate the whole kinome. Proteomics tools have been used to detect the proteins and their phosphorylation levels across tissues in Arabidopsis, maize, and humans [3-5]. To identify the substrates of kinases, functional protein microarrays or peptide arrays were used to probe the human and Arabidopsis kinases *in vitro* [6, 7]. Kinome-wide RNA interference-based functional screening has also been used to interrogate kinase functions with regard to specific biological processes [8, 9]. The functions of kinases have also been investigated by identifying their interactors using large-scale protein interaction networks [10, 11]. Because kinase activities can be affected by small molecules, inhibitor libraries consisting of many small molecules have also been used to screen the kinome to identify potential drug targets [12, 13]. Except for RNAi-based functional screening, these approaches have been mainly focused on the post-transcriptional perspectives to study kinases; for example, proteins and their phosphorylation levels, protein interactors, and kinase inhibitors.

Being proteins themselves, protein kinases are also subjected to transcriptional regulation, and their gene expression patterns can be used to infer their functions. Due to their preferential expression in guard cells among all the MAPK kinases, MPK9 and MPK12 were identified and found to be involved in abscisic acid (ABA) signaling that governs stomatal movement [14]. BUPS1 and BUPS2, two receptor-like kinases (RLKs) that are specifically expressed in pollen tubes, were characterized as regulators for maintaining pollen tube integrity [15]. Two leucine-rich repeat RLKs (LRR-RLKs), MUSTACHES and MUSTACHES-LIKE, were found to have overlapping expression in lateral root primordia and were later identified as regulators of lateral root development [16, 17]. Beyond the individual kinases, co-expression network analysis was conducted to infer the biological processes in which 459 human kinases might participate [18]. These examples illustrate that gene expression analysis can facilitate functional analysis of kinases. However, to our knowledge, such analysis has not yet been conducted on a whole-kinome scale for plant species like Arabidopsis to systematically dissect their functions. With the rapid accumulation of large-scale transcriptome datasets, we expect such analyses will be important to associate Arabidopsis kinases with their functional pathways.

Previously, we developed an approach that we call EXPLICIT to construct a gene expression predictor for Arabidopsis using transcriptome datasets derived from more than 24,545 publicly available RNA-seq runs [19]. The model accurately predicted the expression of 29,182 Arabidopsis genes based on the expression of 1,678 transcription factor (TF) genes. It captured the quantitative relationship between TFs and their target genes and further enabled downstream inference of TF regulators for diverse plant genes and pathways. We envision such an approach can be adapted to analyze the kinome as well.

In this study, we developed a modified approach called EXPLICIT-Kinase to construct a universal Arabidopsis gene expression predictor which accurately predicted the expression of 30,172 non-kinase genes using the expression of 994 protein kinase genes from diverse samples. The model identified the subsets of kinases that best predict the expression of Arabidopsis genes and pathways. Strikingly, many of these predictor kinases are bona fide regulators of the related pathways, as exemplified by those involved in cytokinesis, tissue development, and stress responses. Portions of these predictor kinases also have their orthologues identified as predictor kinases in maize for the corresponding pathways, and they are considered to be conserved predictor kinases between plant species. The conservation between species provide additional evidence to support the novel predictor kinases as bona fide regulators of the pathways involved. Thus, EXPLICIT-Kinase enables a systematic dissection of the functions of the Arabidopsis kinome.

## RESULTS

### Building an Arabidopsis gene expression predictor based on protein kinase genes

In a previous study, we developed the EXPLICIT approach to construct a universal predictor model for accurately predicting Arabidopsis gene expression values based on the expression of TF genes [19]. Here, we developed a modified EXPLICIT-Kinase approach to construct an Arabidopsis gene expression predictor based on protein kinase genes (for simplicity, protein kinases will be referred to as “kinases” hereafter). EXPLICIT-Kinase uses ordinary least squares regression to build a linear predictor model, *Y* = *X B* + *ε*, for expression prediction, with *X* being the log-transformed expression matrix of the kinase genes, *Y* being the log-transformed expression matrix of non-kinase genes, while *B* and ε represent the coefficient matrix and the random errors, respectively (Fig. 1A). The transcriptomes of 26,900 high-quality Arabidopsis RNA-seq runs, available as of April 2020 from the NCBI Sequence Read Archive (SRA) database, were used to train the predictor model. The Arabidopsis genome contains a total of 1,012 kinase genes [2]. After filtering out genes that are expressed at low levels, the model used the expression values of 994 kinase genes to predict the expression of 30,172 non-kinase genes. Similar to the TF-based predictor model, in order to obtain a universal and accurate kinase-based predictor, it was crucial to use a large number of transcriptome samples for model training. Increasing the number of training samples effectively increased the accuracy of the model’s prediction on novel samples, as shown by the correlation (*r*) between the predicted and actual transcriptomes for the test samples (Fig. 1B). Training with 20,000+ samples also reduced the over-fitting within the model to a minimal level, as judged from the difference of *r* between the training and test samples (Fig. 1B). Therefore, all 26,900 RNA-seq transcriptomes were used to train the predictor model.

**Figure 1.**
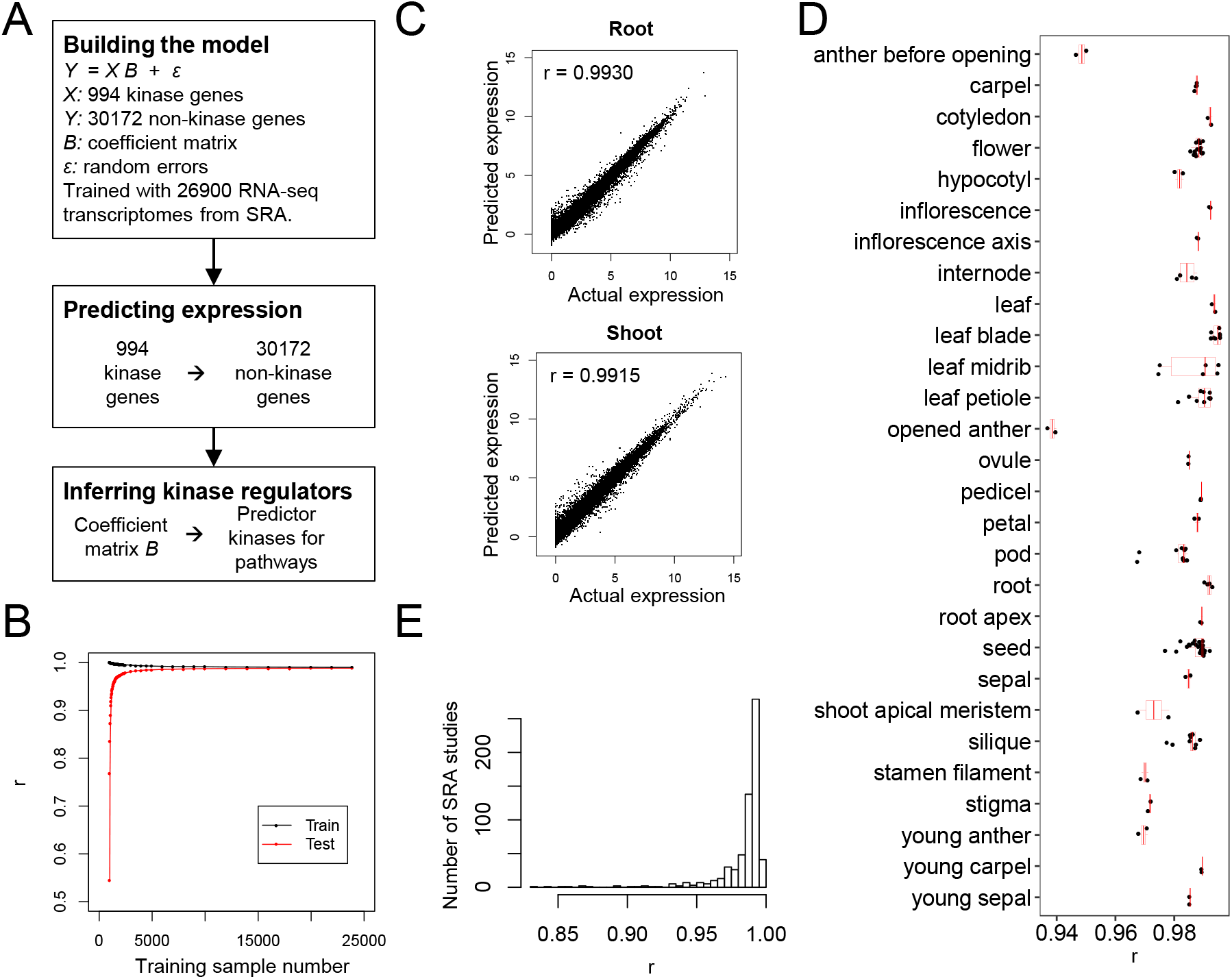
Building a kinase-based Arabidopsis gene expression predictor. **A**. The EXPLICIT-Kinase analysis workflow. **B**. The predicting performance for predictor models trained with different numbers of training samples. Shown are the average Pearson correlation coefficient (*r*) between the predicted and actual transcriptomes for the training samples (black) and independent test samples (red, n=3,000). **C**. The model accurately predicted gene expression in two independent RNA-seq samples from Arabidopsis root and shoot tissues. **D**. The ‘leave-one-out cross-validation’ (LOOCV) test results for the SRA study SRP075604. This study included transcriptome samples from a broad range of Arabidopsis tissues. **E**. LOOCV test results for 630 SRA studies with ≥10 samples. The histogram shows the distribution of each study’s average *r*.

The kinase-based predictor model was then tested to evaluate its performance on transcriptome prediction. Our lab has previously generated two independent Arabidopsis root and shoot RNA-seq transcriptomes to test the TF-based predictor model [19]. We tested the kinase-based model on the same dataset. Using the expression values of 994 kinase genes as input, the model accurately predicted these root and shoot RNA-seq samples, with *r* values of 0.9930 and 0.9915, respectively (Fig. 1C). The accuracy was slightly lower than that of the TF-based predictor model, which had *r* values of 0.9949 and 0.9922 for the root and shoot samples, respectively [19]. The kinase-based model was constructed using 26,900 RNA-seq transcriptomes originating from 1,193 SRA studies, with each study being an independent experiment. We further tested the model on these studies using a ‘leave-one-out-cross-validation’ (LOOCV) strategy. In each LOOCV run, a study was selected and its samples were held out as test samples, while the model was retrained on all other samples and then tested on the held-out samples. The LOOCV results for study SRP075604, which have measured transcriptomes across a broad range of Arabidopsis tissues [20], are shown in Figure 1D. Our kinase-based model showed very good prediction for these samples, with an average *r* of 0.9855. The LOOCV test was further conducted on 630 studies with 10 or more samples. The average *r* for these studies was 0.9836 (Fig. 1E). Thus the kinase-based predictor model achieved very high accuracy on independent test samples, although the accuracy was slightly lower than that of the TF-based model, which had an average *r* of 0.9861 [19].

### Identifying predictor kinases for Arabidopsis genes and gene modules

In our previous EXPLICIT approach, we used the TF-based predictor model to identify predictor TFs for Arabidopsis genes as the subsets of TFs that best predict the expression of the target genes [19]. We also identified predictor TFs for gene modules as those predictor TFs in which the target genes were enriched within the modules. Many of the predictor TFs turned out to be bona fide regulators of the related gene modules.

Following a similar strategy, we used the kinase-based predictor model to identify predictor kinases for Arabidopsis genes. The coefficients within the model were tested for significance. In total, 1,628,911 coefficients with *p*-value ≤1E-09 (Bonferroni-corrected *p*-value ≈0.05) were identified in which the values were significantly different from zero (Table S1). Based on these coefficients, significantly interacting kinase–non-kinase gene pairs were retrieved, and the kinases connected to a specific non-kinase gene by these gene pairs were considered to be the predictor kinases of that gene. A gene’s predictor kinases constitute the subset of kinases for which the expression best predicts that gene’s expression value. On average, each gene has 54 predictor kinases. Beyond individual genes, predictor kinases were also identified for gene modules. A kinase was considered to be a predictor kinase for a gene module if it was shared by the genes within the module as a common predictor kinase with enrichment *p*-value ≤1E-05 (adjusted to decrease the false discovery rate using the Benjamini-Hochberg procedure).

Previously, we constructed an Arabidopsis gene co-expression network based on the graphical Gaussian model (GGM) and identified 1,085 gene co-expression modules from the network [19]. Here we expanded the modules by adding the outside genes that were connected to four or more genes within the modules, reasoning that they should have similar expression patterns to the modules (Table S2). Gene Ontology enrichment analysis indicated that these expanded modules participate in a broad range of biological processes (Table S3). We then identified predictor kinases for these modules (Table S4). Strikingly, the identified predictor kinases are usually the bona fide kinase regulators for the pathways associated with the modules, as demonstrated by the examples discussed below.

### Predictor kinases identified for cytokinesis and related processes

The cell cycle and cytokinesis, pivotal processes for all eukaryotic species, involve multiple steps that require precise coordination and regulation. As modulators of protein activities, protein kinases are important regulators of the cell cycle and cytokinesis. Among the modules identified from the GGM co-expression network, Module #4 was enriched with 64 *cell cycle* genes (*p* = 1.79E-58) and is thus considered to be a module for the cell cycle and cytokinesis (Fig. 2A). Using EXPLICIT-Kinase, we identified 28 predictor kinases for the module (Fig. 2B). Among them are five cyclin dependent kinases (CDKB1;1, CDKB1;2, CDKB2;1, CDKB2;2, and CDKD1;1) that control cell cycle progression [21-23], three aurora kinases that regulate chromosomal segregation (AUR3) and formation of the division plane orientation during cytokinesis (AUR1, AUR2) [24, 25], and two mitogen-activated protein kinase kinase kinases (MAPKKK) (ANP2 and ANP3) that positively regulate cell division and growth [26]. Also identified were RUNKEL and TIO, two kinases required for proper phragmoplast organization and expansion during cytokinesis [27, 28]. Indeed, of the 28 predictor kinases, 16 (57.1%) of them are known regulators of the cell cycle or cytokinesis (see Table S5 for detailed references, same as below). We checked the expression patterns of the uncharacterized predictor kinases across Arabidopsis tissues using the transcriptomes from the AtGenExpress project [29]. The expression of these predictor kinase genes is similar to that of the genes within Module #4, with relatively high expression levels in the tissues undergoing active cell divisions such as the shoot apex (Fig. S1), implying that they might also be regulators of the cell cycle.

**Figure 2.**
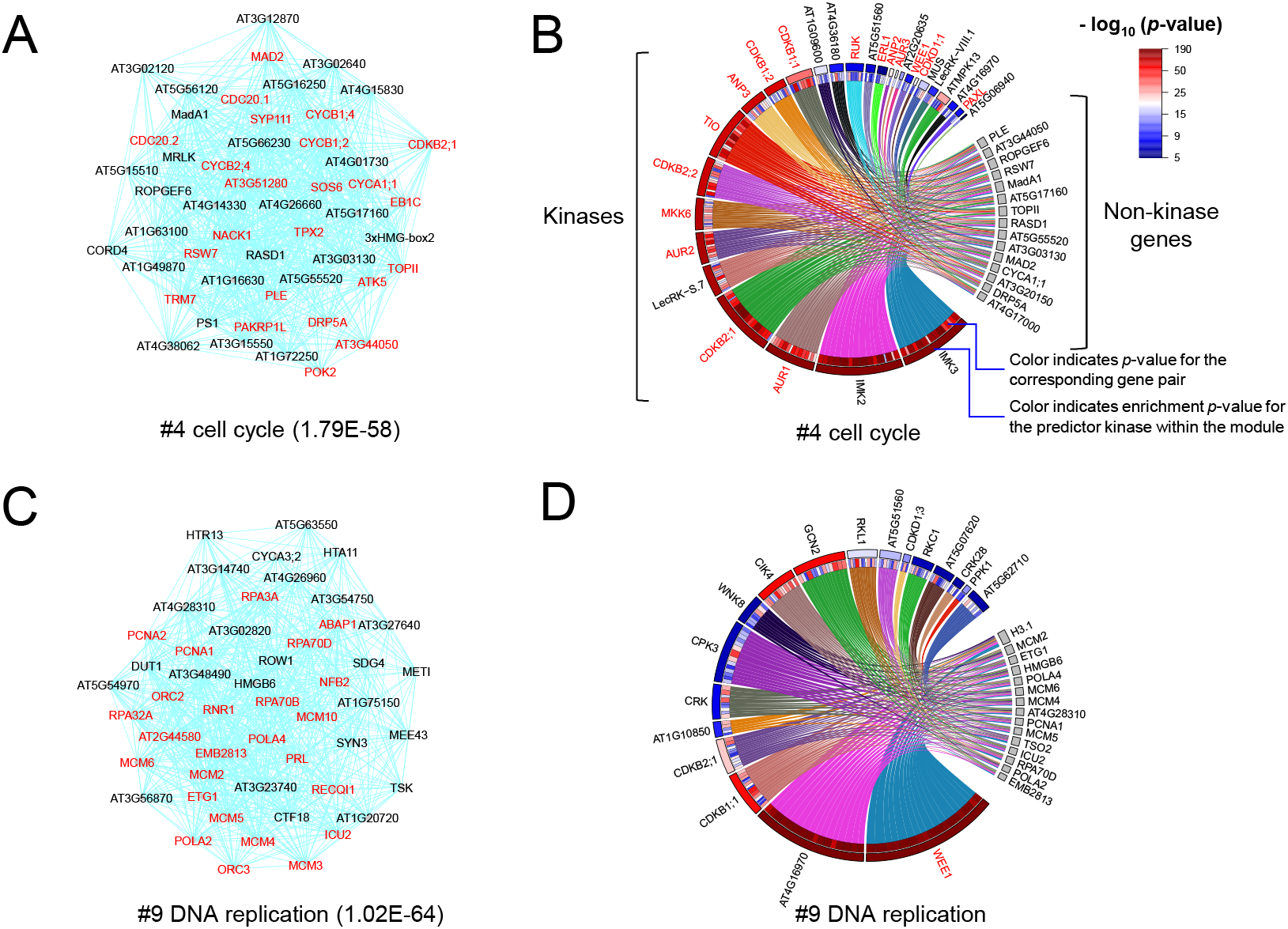
Predictor kinases identified for gene modules that function in cytokinesis and DNA replication. **A, C**. The gene co-expression modules for cytokinesis (#4) and DNA replication (#9). The module ID and its enriched GO term (and its *p*-value) are shown. Gene names shown in red text are the genes possessing the GO term. Only the top 50 genes with the highest number of connections within the modules are shown due to space limitation. **B, D**. Predictor kinases identified for the modules. Chord diagrams connecting the modules’ predictor kinases to the genes whose expression values they significantly predict are shown. The links are grouped by predictor kinases. *p*-values for the predictor kinase–non-kinase gene pairs and the enrichment of the predictor kinases are indicated by colors. Gene names in red text are known kinase regulators of the pathways involved.

In addition to the cell cycle, we also identified 18 predictor kinases for a related module functioning in *DNA replication* (#9, *p* = 1.02E-64) (Fig. 2C, D). The highest ranking predictor kinase is WEE1, which functions as a DNA replication checkpoint kinase to ensure genome integrity [30]. The module shares a handful of predictor kinases with the cytokinesis module, such as CDKB1;1 and CDKB2;1, but the roles of other predictor kinases in DNA replication remain to be investigated.

### Predictor kinases identified in tissue development

As a multicellular organism, the development of Arabidopsis tissues and organs involves signal communication and coordination between different cells, during which kinases play important roles. EXPLICIT-Kinase enabled the identification of the kinases that function in these processes. As an example, Module #1 from the GGM network was considered a module for pollen tube development, since it was enriched with 27 *pollen tube development* genes (*p* = 1.59E-20) (Fig. 3A). Previously, we identified 80 predictor TFs for the module, and 22 of them were known regulators of pollen tube development [19]. Here, using EXPLICIT-Kinase, we identified 64 predictor kinases for the module (Fig. 3B). The large number of predictor kinases was consistent with the observation that many kinases play roles during pollen tube development. Among the identified predictor kinases, BUPS1 and BUPS2 are two receptor-like kinases (RLKs) that regulate pollen tube integrity and sperm release [15]; PRK1, PRK3, PRK6, and PRK8 are tip-localized RLKs that control pollen tube growth [31]; LIP1 and LIP2 are tip-anchored RLKs involved in guiding pollen tube growth into the micropyle [32]; PERK5 and PERK12 are two Proline-rich Extension-like receptor kinases (PERKs) that modulate pollen tube polar growth [18]; and CPK17 and CPK34 are calcium-dependent protein kinases (CDPKs) that are also required for polarized tip growth [33]. Indeed, 22 out of the 64 predictor kinases are known regulators of pollen tube growth or pollen tube functions (Table S5). Among these 22 known kinase regulators, 16 are RLKs and five are CDPKs or calcineurin B-like (CBL)-interacting protein kinases (CIPKs), and all are the type of kinases involved in signaling transduction and relaying. Interestingly, among the 42 uncharacterized kinases, 29 are also RLKs, including four PERKs (PERK4, PERK6, PERK7, and PERK11). According to AtGenExpress and another transcriptome analysis of pollen tubes [29, 34], most of the uncharacterized predictor kinases are also specifically or highly expressed in pollen tubes, indicating their possible involvement in pollen tube development or functions (Fig. S2).

**Figure 3.**
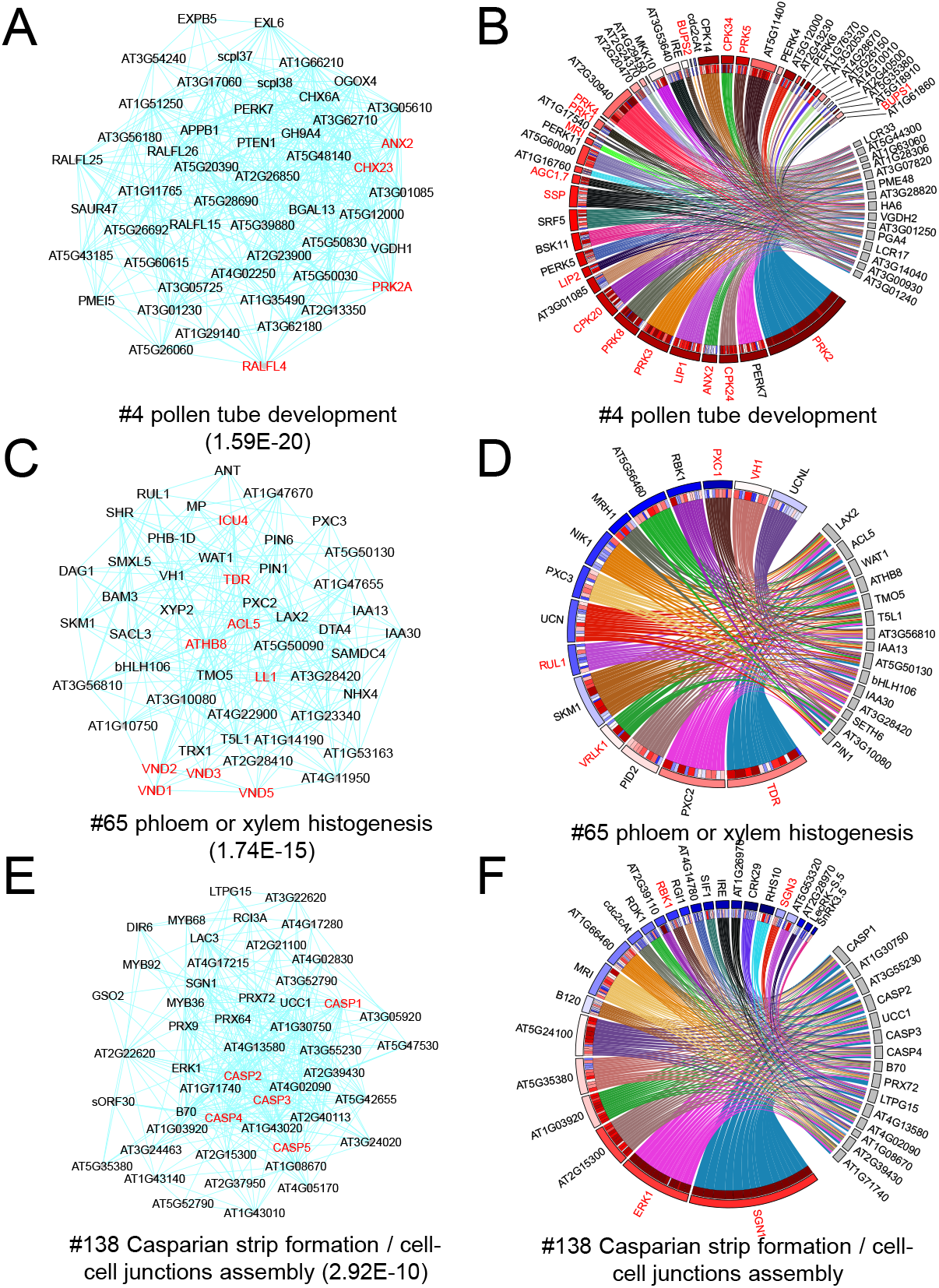
Predictor kinases identified for gene modules involved in tissue development or functions. **A, C, E**. The gene co-expression modules for pollen tube development (#4), vascular development (#65), and Casparian strip formation (#138). **B, D, F**. Predictor kinases identified for the four modules. Only a portion of the predictor kinases are shown for Module #4 **(A)** due to space limitations. Legends are the same as in Figure 2.

We also identified 15 predictor kinases for Module #65 that functions in vascular tissue development, which was enriched with 12 genes related to *phloem or xylem histogenesis* (*p* = 1.74E-15) (Fig. 3C, D). The highest ranking predictor kinase is TDR, a leucine-rich repeat RLK (LRR-RLK) that mediates cellular communication between vascular stem cells and governs xylem-phloem patterning [35]. Also identified are four other LRR-RLKs important for vascular or secondary cell wall development: RUL1, PXC1, VH1, and VRLK1 [36-39]. Thus, five out of the 15 predictor kinases are known regulators of vascular development. Among the 10 uncharacterized kinases, four are also LRR-RLKs, such as PXC2 and PXC3 that are also expressed in vascular tissues [39], and three, including PID2, UCN, and UCNL, are AGC (cAMP-dependent, cGMP-dependent and protein kinase C) family kinases. Interestingly, another AGC family kinase, PAX, was also enriched in the module, but its *p*-value (5.68E-05) was slightly higher than our selected cutoff. Nevertheless, PAX is a kinase that regulates auxin flux and protophloem development [40], indicating that the other three AGC kinases might also have similar roles during vascular development.

As a comparison to the vascular module, we identified 25 predictor kinases for Module #138 that functions in Casparian strip and endodermis formation in the roots; the module contained five genes that encode Casparian strip domain proteins (*CASP1*-*5*) (Fig. 3E, F). The highest ranking predictor kinase was Schengen1 (SGN1) that regulates Casparian strip integrity and positioning [41]. Also identified were Schengen3, a LRR-RLK involved in endodermis integrity, and ERK1 and RBK1, another two kinases recently characterized as being required for Casparian strip formation [42, 43]. Interestingly, among the other 21 uncharacterized predictor kinases, seven were also LRR-RLKs, but their possible involvement in Casparian strip development remains to be investigated.

Additionally, predictor kinases were also identified for other development-related modules, such as those for the development of root hairs (#5), anthers (#78), pollen/gametophyte (#60), the phyllome (#198), floral organs (#132), the shoot apical meristem (#197), the root meristem (#103), those that regulate cell wall loosening/cell growth (#164) and fertilization (#245), as well as those involved in the biosynthesis of the primary cell wall (#82) and secondary cell wall (#11) (Table S4). These predictor kinases could help to enhance our understanding of the roles played by kinase regulators in various developmental processes.

### Predictor kinases identified for response to stresses and environmental stimuli

As sessile organisms, plants face environmental stresses constantly. They rely on efficient signaling networks to respond to stresses for their survival. Protein kinases are important components of plant signaling networks. We sought to identify the kinases that might function in the stress response. In our previous study, we have identified GO regulons from the GGM gene co-expression network [19]. A GO regulon contains genes from the same GO term that are also connected as a single component within the GGM gene co-expression network, and thus it can be considered to be a gene co-expression module for that GO term. We selected the GO regulons related to the response to stresses or environmental stimuli and inferred their predictor kinases.

As an example, the regulon for *response to wounding* (GO:0009611) contains 90 genes (Fig. 4A). We identified 34 predictor kinases for the regulon, including five known regulators of the wounding stress response: PEPR1, PEPR2, MPK3, OXI1, and P2K1 (Fig. 4B and Table S6) [44-47]. According to AtGenExpress [48], among the other predictor kinases, 15 are also induced by wounding treatment in Arabidopsis leaves, such as SIF4, MPKKK14, and LYK5 (Fig. S3), indicating their possible involvement in the wound response. We also analyzed the GO regulons for the *response to bacterium* (GO:0009617), *cold* (GO:0009409), *fungus* (GO:0009620), *abscisic acid* (GO:0009737), *jasmonic acid* (GO:0009753), *osmotic stress* (GO:0006970), *oxidative stress* (GO:0006979), and *water deprivation* (GO:0009414), and identified between 31 and 119 predictor kinases for each of them (Table S6).

**Figure 4.**
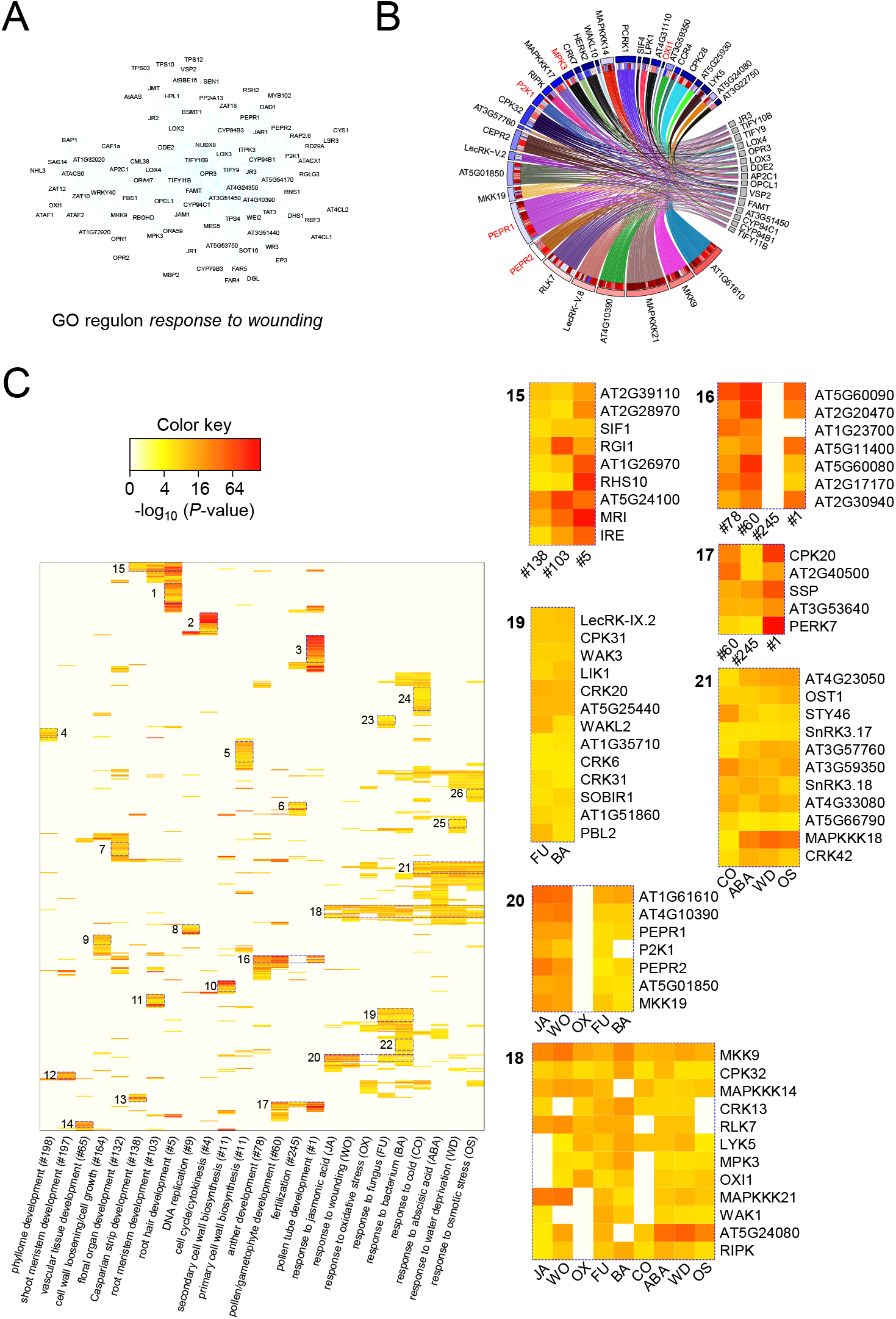
Predictor kinases identified for the GO regulons involved in stress responses. **A**. A GO regulon for the *response to wounding* (GO:0009611). All genes within this regulon possess the GO term GO:0009611, and they are connected as a single component within the GGM gene co-expression network. **B**. Predictor kinases identified for the regulon GO:0009611. **C**. A heatmap showing the shared and distinct predictor kinases among the gene modules and regulons. On the left is an overall heatmap showing the *p*-values for 529 predictor kinases (rows) for the modules or regulons (columns), with areas of interest shown in 26 boxes. On the right are selected boxes containing shared predictor kinases. See Figure S4 for a more detailed heatmap.

A comparison between the stress response regulons, as well as between the modules for cytokinesis and development, revealed distinct and shared predictor kinases (Fig. 4C and Fig. S4). All these modules and regulons had a total of 552 predictor kinases. In general, compared to the stress response modules, the cytokinesis and development-related modules have more unique predictor kinases, which are not shared with other modules or regulons. Most of the cytokinesis and development-related modules have a fair number of unique predictor kinases (Boxes 1 – 14 in Fig. 4C). The specificities of these predictor kinases imply their specific functions in cytokinesis or in development of the corresponding tissues. At the same time, some related tissues share portions of the common predictor kinases, such as those shared between the root meristem, root hairs, and the Casparian strip (Box 15, Fig. 4C), those shared between anthers, the gametophyte, and pollen tubes (Box 16, Fig. 4C), and those shared between the gametophyte, egg cells, and pollen tubes (Box 17, Fig. 4C).

For the stress response regulons, we observed shared as well as distinct predictor kinases between the different regulons, and these can be considered to be cross-talk and specificities among the different stress response pathways. A handful of predictor kinases were shared by more than five different stresses, such as OXI1, RIPK, MPK3, MKK9, MPKKK14, and LYK5 (Box 18, Fig. 4C). Among them, OXI1, RIPK, and MPK3 have been shown to regulate reactive oxygen species (ROS) signaling, highlighting the importance of ROS in the stress response [45, 49]. The responses to bacteria and fungi shared two groups of common predictor kinases: the first group (Box 19, Fig. 4C) is specific to the responses to bacteria and fungi, with kinases like SOBIR1, PBL2, and LIK1; the second group (Box 20, Fig. 4C) is also shared with the response to wounding stress and jasmonic acid, and contains kinases like PEPR1 and PEPR2. The responses to water deprivation, cold, osmotic stress, and abscisic acid (ABA) also shared a group of common predictor kinases (Box 21, Fig. 4C), which included SnRK3.17, MAPKKK18, and OST1. Among these, MAPKKK18 is an ABA-responsive kinase involved in the drought stress response, while OST1 also functions in ABA-dependent pathways to module the responses to cold, drought, and osmotic stresses [50-53]. Additionally, predictor kinases specific to the responses to bacteria, fungi, cold, water deprivation, and osmotic stress were also identified (Boxes 22–26 in Fig. 4C). Thus, our analysis identified the individual kinases and placed them into the potential pathways in which they might participate.

### Conserved predictor kinases between Arabidopsis and maize

Our analysis identified many novel predictor kinases for a broad range of gene modules and pathways. We asked whether these novel predictor kinases are conserved between different plant species, in this case *Arabidopsis thaliana* (a dicot) and maize (*Zea mays*, a monocot). If a kinase in Arabidopsis and its orthologue in maize are both predictor kinases for the same pathway in both species, the predictor kinase is conserved, which provides additional evidence to support the kinase as being a bona fide regulator of the pathway involved. Therefore, we collected large-scale maize transcriptome data, identified gene co-expression modules via GGM network analysis (Table S7), and inferred predictor kinases for the modules (Table S8). The maize model had only 12,959 RNA-seq runs as training samples and identified only 399,996 significant predictor kinase– non-kinase gene pairs. Therefore, fewer predictor kinases were identified for maize genes and pathways than in Arabidopsis. However, a comparison between the Arabidopsis and maize modules did reveal conserved predictor kinases, as demonstrated by the following examples.

The maize module Zm#3 contained 87 genes with orthologues in the Arabidopsis cytokinesis module (#4), including genes like *cyc1, cyc3, cyc5, cyc6, cyc7*, and *cyc8* that encode cyclin family proteins. Thus, Zm#3 was considered to be a cytokinesis module for maize. Using the maize kinase-based predictor model, we identified 27 predictor kinases for the module (Fig. 5A). Ten of orthologues of these predictor kinases are among the known cytokinesis regulators that were also identified from the Arabidopsis module; examples are Zm00001d008815 (AUR1), Zm00001d034164 (AUR3), Zm00001d044672 (CDKB1;1), and Zm00001d050329 (ANP1) (the Arabidopsis orthologues of the maize kinases are shown in parentheses, same as below). Interestingly, we also found eight novel maize predictor kinases which have their orthologues among the novel predictor kinases for cytokinesis in Arabidopsis, such as Zm00001d052323 (IMK2), Zm00001d028796 (IMK2), Zm00001d037371 (LecRK-S.7), Zm00001d035476 (LecRK-VIII.1), Zm00001d000034 (AT4G16970), Zm00001d045445 (AT5G51560), Zm00001d002599 (AT4G36180), and Zm00001d026063 (AT4G36180). These novel predictor kinases were identified from the cytokinesis modules from both Arabidopsis and maize, indicating they might have conserved functions in cell cycle regulation in both species. Similarly, the maize module Zm#22 was considered to be a module for DNA replication, and it shares 73 orthologous genes with its Arabidopsis counterpart (Module #9). Ten predictor kinases were identified from the maize module, and the top three are conserved, including Zm00001d000034 (AT4G16970), Zm00001d053998 (WEE1), and Zm00001d044672 (CDKB1;1) (Fig. 5B). Among them, the novel predictor kinases Zm00001d000034 (AT4G16970) were ranked as the top first and second predictor kinases in maize (*p* = 1.48E-281) and Arabidopsis (*p* = 2.61E-190), respectively. Thus, they can be considered to be ideal candidate regulators for DNA replication.

**Figure 5.**
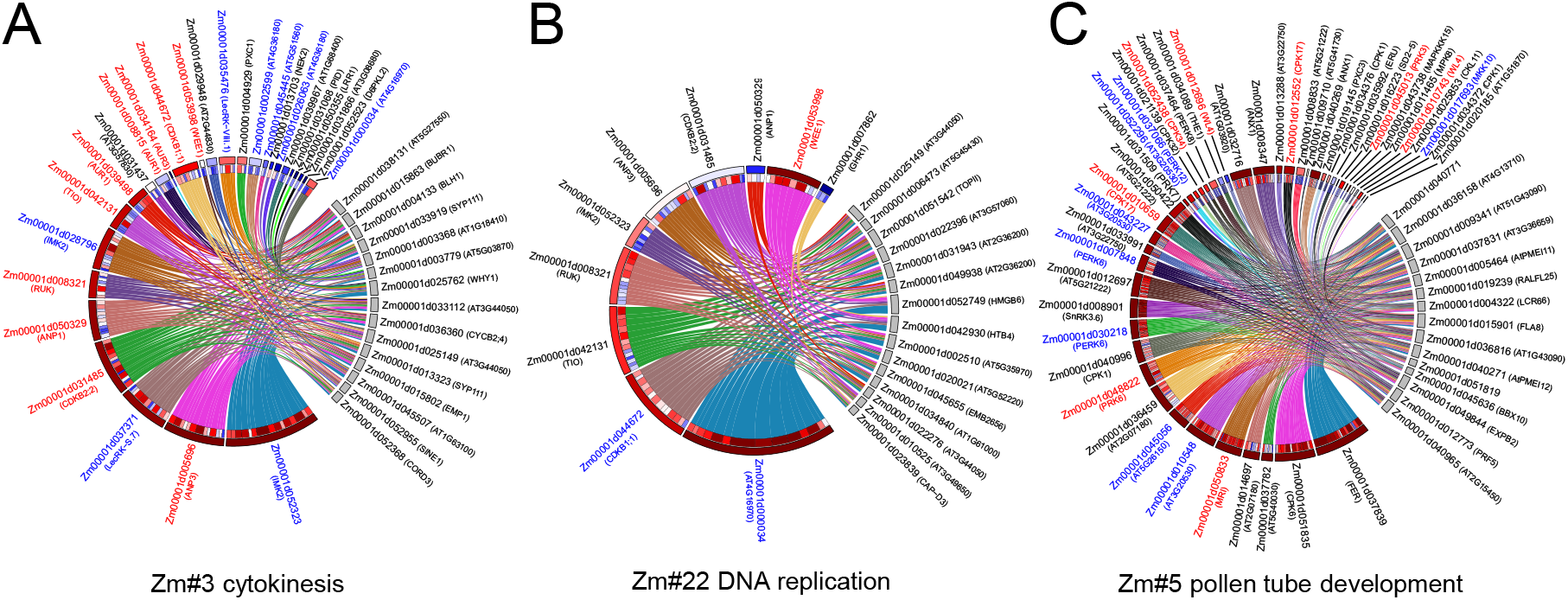
Predictor kinases identified for the maize gene modules. The modules involved in cytokinesis **(A)**, DNA replication **(B)**, and pollen tube development **(C)** are shown. The Arabidopsis orthologues of the maize genes are shown in parentheses. Names in red and blue are the known and novel conserved predictor kinases, respectively, identified from both maize and Arabidopsis gene modules of the same pathways. All other legends are the same as in Figure 2.

As another example, the maize module Zm#5 shared 43 orthologous genes with Arabidopsis module #1, and it is considered to be a module for pollen tube development. We identified 63 predictor kinases for the module, including 20 conserved kinases shared with Arabidopsis Module #1 (Fig. 5C). Among these 20 conserved predictor kinases, nine are known regulators of pollen tube development in Arabidopsis, while the other 11 are novel predictor kinases, and their roles will require further investigation; examples are Zm00001d030218 (PERK6), Zm00001d017693 (MKK10), and Zm00001d010548 (AT3G20530). Interestingly, a known Arabidopsis pollen tube regulator kinase called FER [54], an orthologue of Zm00001d037839, was only identified as a predictor kinase in maize but not in Arabidopsis.

Thus, our comparison analysis identified both known and novel conserved predictor kinases between Arabidopsis and maize. These kinases might have conserved functions between species, and the novel ones can be considered as prioritized candidates for future functional studies.

## DISCUSSION

In the current study, we developed the EXPLICIT-Kinase approach to build a model to predict Arabidopsis transcriptomes based on the expression of 994 protein kinase genes. The approach was modified from our previous EXPLICIT approach which used the expression of TF genes to predict Arabidopsis transcriptomes [19]. Similar to the original EXPLICIT, after training with more than 25,000 samples, EXPLICIT-Kinase generated a highly accurate predictor model that can faithfully reconstitute Arabidopsis transcriptomes from a broad range of tissues as well as from independent SRA studies (Fig. 1B-E). More importantly, for each non-kinase gene, the model identified a subset of kinases with expression values that best predict the expression of the non-kinase gene. We called this subset of kinases the ‘predictor kinases’. Being proteins themselves, protein kinase genes are subjected to transcriptional regulation, and their expression patterns provide important clues about their functions. The assignment of predictor kinases for non-kinase genes effectively summarizes the expression patterns of kinase genes, which can then be used to infer predictor kinases for gene modules via enrichment assays. The predictor kinases for gene modules associate the kinases with the biological pathways they participate in, providing a way to systematically dissect the functions of the Arabidopsis kinome.

The EXPLICIT-Kinase approach identified predictor kinases for gene modules involved in a broad range of biological processes, and the predictor kinases were often the bona fide kinase regulators for the related pathways, as exemplified by the examples discussed above. In plants, kinases regulate many aspects of life actives, including cytokinesis, tissue development, and responses to environmental stresses and stimuli. Our approach identified predictor kinases for gene modules functioning in all these aspects. For example, 28 predictor kinases were identified for the cytokinesis module, 15 of which are known regulators (Fig. 2A, B); 64 and 15 predictor kinases were identified for modules involved in pollen tube and vascular tissue development, respectively, among which 19 and five are known regulators (Fig. 3A-D); and 34 predictor kinases were identified for the response to wounding, among which five are known regulators (Fig. 4B). Within these modules, as well as those not discussed in detail, there were many uncharacterized and novel predictor kinases. These novel predictor kinases usually have expression patterns that are similar to those of the genes within the modules (Fig. S1-3), implying their involvement in the corresponding pathways. Further support comes from the functional conservation of predictor kinases in both Arabidopsis and maize, such as IMK2, IMK3, and LecRK-S.7 for the cytokinesis module, as well as AT4G16970 for the DNA replication modules. However, it should be noted that there are many fewer publicly-available maize RNA-seq samples than there are in Arabidopsis, which might limit attempts to identify additional conserved predictor kinases. We expect that, with more RNA-seq samples for maize and other plant species becoming available in the future, more conserved predictor kinases will be identified.

Our approach predicts the functions of Arabidopsis kinases and associates them with the pathways they might participate in based on their expression patterns. The results showed that some gene modules contained multiple predictor kinases from the same kinase family, indicating functional redundancy among them. For example, six PERKs were identified as predictor kinases for pollen tube development (Module #5), nine LRR-RLKs and three AGC kinases were identified as predictor kinases for vascular development (#65), and seven LRR-RLK were identified as predictor kinases for Casparian strip development (#138). In a previous study, Wu and colleagues systematically investigated the genome-wide expression patterns of 223 LRR-RLK genes [16]. They noted that only a limited number of LRR-RLKs generated visible phenotypes when knocked out, possibly due to gene redundancy. Our analysis revealed such redundancy at the expression level and provided a guidance to generate multiple knock-out mutants to further investigate the functions of protein kinases. However, completely elucidating the function of a kinome requires identifying the substrates for the protein kinases. It should be noted that our EXPLICIT-Kinase approach is not designed to identify substrates for protein kinases, although it is possible that kinase genes and their substrate genes might be co-expressed. Thus, EXPLICIT-Kinase should be used in conjunction with other systems biology approaches, such as protein interaction networks, protein microarrays, proteomics tools, as well as large-scale kinase mutants, to fully investigate the functions of the Arabidopsis kinome.

On another note, our previous and current studies on EXPLICIT and EXPLICIT-Kinase have demonstrated that these two approaches work well to dissect the functions of TFs and protein kinases. We were able to build Arabidopsis gene expression predictors by using either TFs or protein kinase genes as regressors for model construction. The models further enabled downstream inference of predictor TFs or predictor kinases for genes and pathways, and they were often found to be the bona fide regulators of the corresponding pathways. We expect that similar models can also be used to dissect the functions of other protein categories, such as all transporters or all protein ligases, for both Arabidopsis and other plant and animal species.

In conclusion, we have developed a computational approach that we call EXPLICIT-Kinase to dissect the functions of the Arabidopsis kinome based on their expression patterns. This approach can be used to guide further investigations on Arabidopsis protein kinases. We also developed a companion software package, called EXPLICIT-Kinase, which is freely available through GitHub (https://github.com/MaShisongLab/explicit-kinase).

## MATERIAL AND METHODS

### Building the kinase-based expression predictor

We developed the EXPLICIT-Kinase approach to build a kinase-based expression predictor model for Arabidopsis. EXPLICIT-Kinase was modified from our previous EXPLICIT approach [19]. The two approaches are almost identical, except that while EXPLICIT used TF genes as regressors to build the predictor model, EXPLICIT-Kinase uses kinase genes instead. Briefly, publicly available Arabidopsis RNA-seq raw data was downloaded from the NCBI SRA database as of April 2020 and processed as previously described [19]. After filtering out low-quality, single-cell, or small RNA-seq samples, the transcriptomes from 26,900 high-quality bulk RNA-seq runs were retained. Their gene expression values (count per millions, CPM) were log-transformed via log_2_(CPM + 1) and merged into a large expression matrix.

A full list of all 1,012 Arabidopsis protein kinases were obtained from a previous study on plant kinomes [2]. According to the list, and after filtering out genes expressed at low levels, two expression matrices, *X* for 994 kinase genes and *Y* for 30,172 non-kinase genes, were extracted. These matrices were used to train a linear model, *Y* = *X B* + *ε*, for expression prediction, with *B* and ε representing the coefficient matrix and the random errors, respectively. The coefficient matrix *B* was estimated using ordinary least squares regression. The resulting *B* was then used for gene expression prediction via the formula: *Y*_*p*_ = *X*_*t*_ *B*, with *X*_*t*_ being the kinase expression matrix for the testing samples, and *Y*_*p*_ being the predicted expression matrix for non-kinase genes. The Pearson correlation coefficients (*r*) between the predicted and actual transcriptomes were then calculated to evaluate the model’s predicting performance.

Two independent RNA-seq transcriptomes generated previously from Arabidopsis root and shoot samples in our lab were used to test the predictor’s predicting performance [19]. Additionally, the predictor was also evaluated via a LOOCV (‘leave-one-out cross-validation’) strategy. In each LOOCV run, we held out the samples from a single SRA study, retrained the model using all other samples, and then tested the retrained model on the held-out samples.

### Identifying predictor kinases for genes and gene modules

Our previous EXPLICIT approach was developed to identify predictor TFs for genes and gene modules [19]. EXPLICIT-Kinase uses the same procedure to identify predictor kinases. Briefly, after training the kinase-based model with all 26,900 RNA-seq transcriptomes, the individual coefficient of the coefficient matrix *B* was checked for significance. The coefficients with *p*-values ≤1E-09 were considered to be significant, and their values were significantly different from zero. The kinase-non-kinase gene pairs associated with these coefficients were extracted, and the kinases connected to a non-kinase gene via these gene pairs were considered to be the predictor kinases for that non-kinase gene.

Beyond individual genes, predictor kinases were also identified for gene modules via enrichment assays based on the hypergeometric distribution. A kinase was considered to be a predictor kinase for a gene module if it was shared by multiple genes within the module as a common predictor kinase with enrichment. The enrichment *p*-value was calculated as:

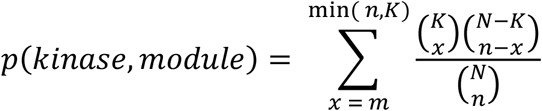

Where *N, K, n*, and *m* represent the total number of non-kinase genes within the genome (*N*), the total number of non-kinase genes that have that kinase as a predictor kinase within the genome (*K*), the number of non-kinase genes within the module (*n*), and the number of non-kinase genes having that kinase as a predictor kinase within the module (*m*). The raw *p*-values were adjusted for multiple testing via the Benjamini-Hochberg procedure after conducting the analysis for all kinases [55]. The kinases with adjusted *p*-value ≤1E-5 were considered to be predictor kinases for the gene module. Chord diagrams were drawn to visualize the interactions between the predictor kinases and the genes within a module using the ‘circlize’ package in R [56]. A software package, also called EXPLICIT-Kinase, was developed to conduct the analysis.

We then identified predictor kinases for Arabidopsis gene modules and GO regulons. The modules were obtained from our previous Arabidopsis GGM gene co-expression network analysis [19]. We expanded the original modules by adding the outside genes that were connected to four or more genes within the modules. Similarly, the GO regulons were also retrieved from our previous analysis [19]. Briefly, a GO-specific subnetwork was extracted from the GGM gene co-expression network by using all the genes possessing the same GO BP (‘biological process’) term (experimentally validated) as nodes. The subnetwork was then analyzed to obtain its largest connected component, which was then treated as a GO regulon.

### Comparative analysis between Arabidopsis and maize

We downloaded and processed 12,959 high-quality maize RNA-seq transcriptomes from NABI SRA. After filtering out genes expressed at low levels, these transcriptomes were used to construct a maize GGM gene co-expression network following a procedure described previously [57]. The network was clustered via the MCL clustering algorithm to identify gene co-expression modules [58]. Orthologous gene pairs between maize and Arabidopsis were identified via AHRD [59] (https://github.com/groupschoof/AHRD). The maize genes are considered as orthologues to the Arabidopsis genes which share the highest homology as identified by AHRD. Conserved gene modules between maize and Arabidopsis were identified as those modules that shared multiple orthologous genes. Using a maize protein kinases list [2], a kinase-based predictor model was developed for maize. The predictor kinases for the conserved gene modules were compared to identify conserved predictor kinases between Arabidopsis and maize.

## SUPPLEMENTAL FIGURES AND TABLES

**Figure S1**. Tissue-specific expression patterns for the predictor kinases and the genes of the cytokinesis module.

**Figure S2**. Tissue-specific expression patterns for the predictor kinases of the pollen tube development module.

**Figure S3**. Gene regulation pattern upon wounding stress treatment for the predictor kinases of the wounding stress response GO regulon.

**Figure S4**. A detailed heatmap showing the shared and distinct predictor kinases among the gene modules and regulons.

**Table S1**. The significantly interacting kinase–non-kinase gene pairs.

**Table S2**. The genes contained within the modules from the Arabidopsis GGM gene co-expression network.

**Table S3**. Gene Ontology enrichment analysis results for the Arabidopsis GGM gene co-expression modules.

**Table S4**. Predictor kinases identified for the Arabidopsis GGM gene co-expression modules.

**Table S5**. References for the known predictor kinases identified for the Arabidopsis GGM gene co-expression modules.

**Table S6**. Predictor kinases identified for the Arabidopsis GO regulons.

**Table S7**. The genes contained within the modules from the maize GGM gene co-expression network.

**Table S8**. Predictor kinases identified for the maize GGM gene co-expression modules.

## ACKNOWLEDGEMENT

We thank USTC Supercomputing Center and USTC School of Life Sciences Bioinformatics Center for providing the computing resources. This work was supported by grants from the National Natural Science Foundation of China (31770268), the Strategic Priority Research Program of the Chinese Academy of Sciences (XDA24010303), the Fundamental Research Funds for the Central Universities (WK2070000091), and University of Science and Technology of China (Start-up fund to S.M.).

