## Supplementary figures and images for "EXPLICIT-Kinase: a gene expression predictor for dissecting the functions of the Arabidopsis kinome"

### Fig. S1

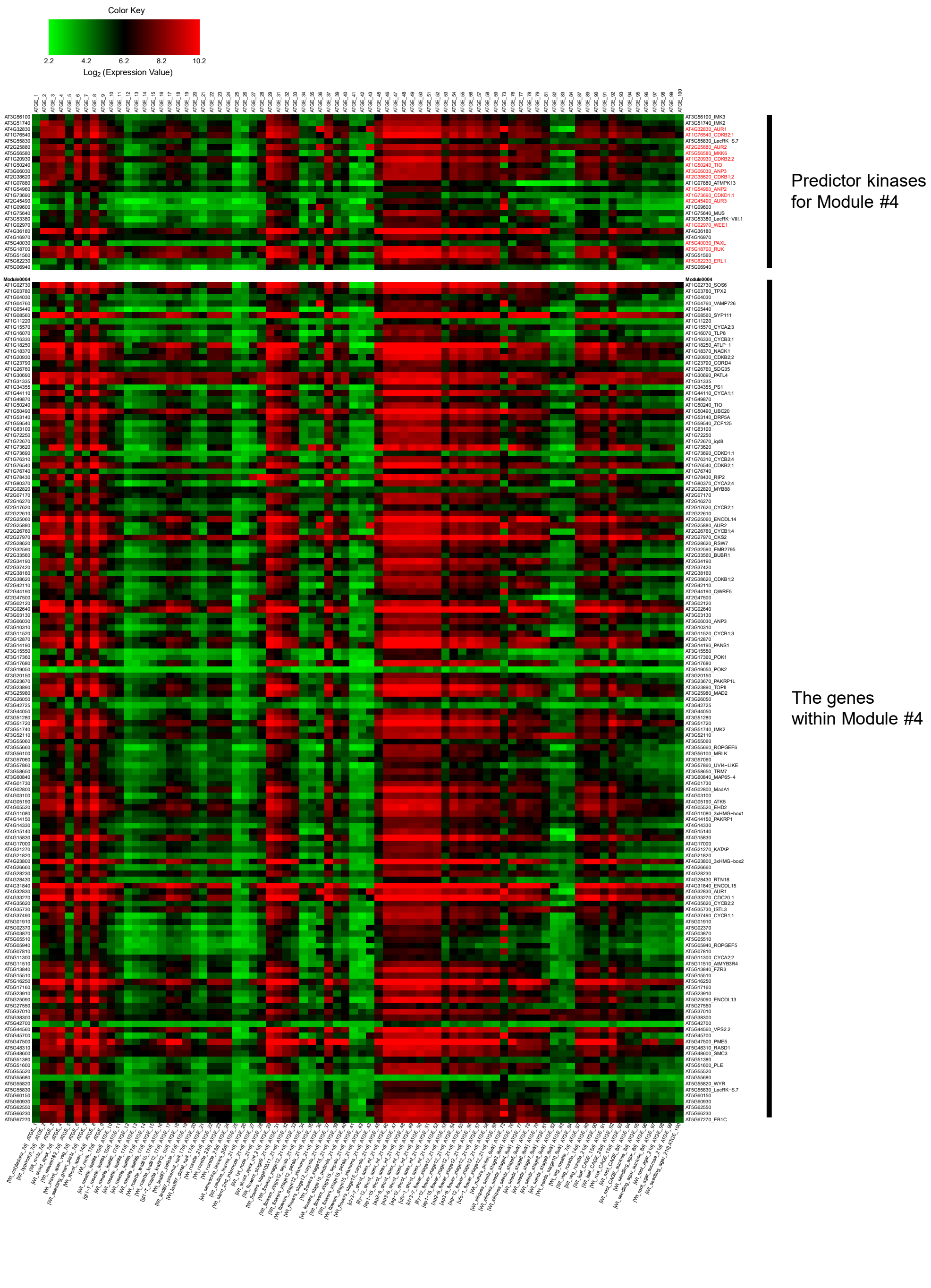

### Fig. S2

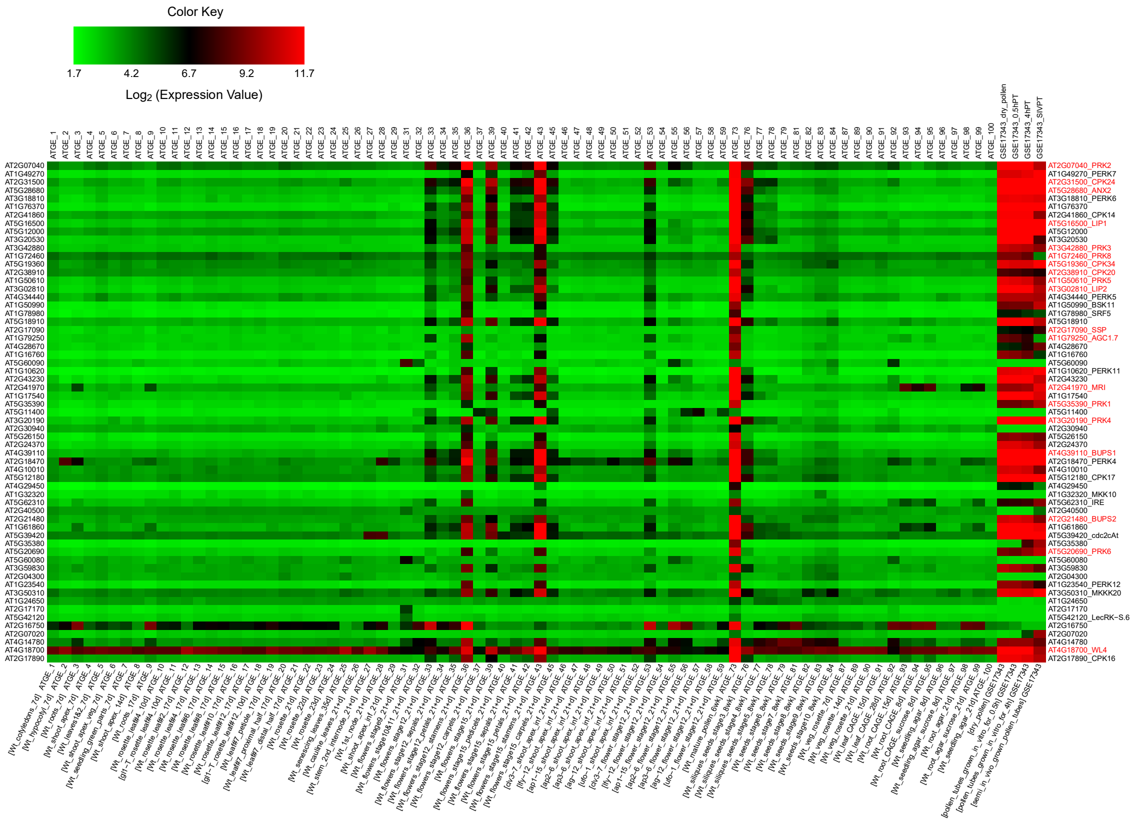

### Fig. S3

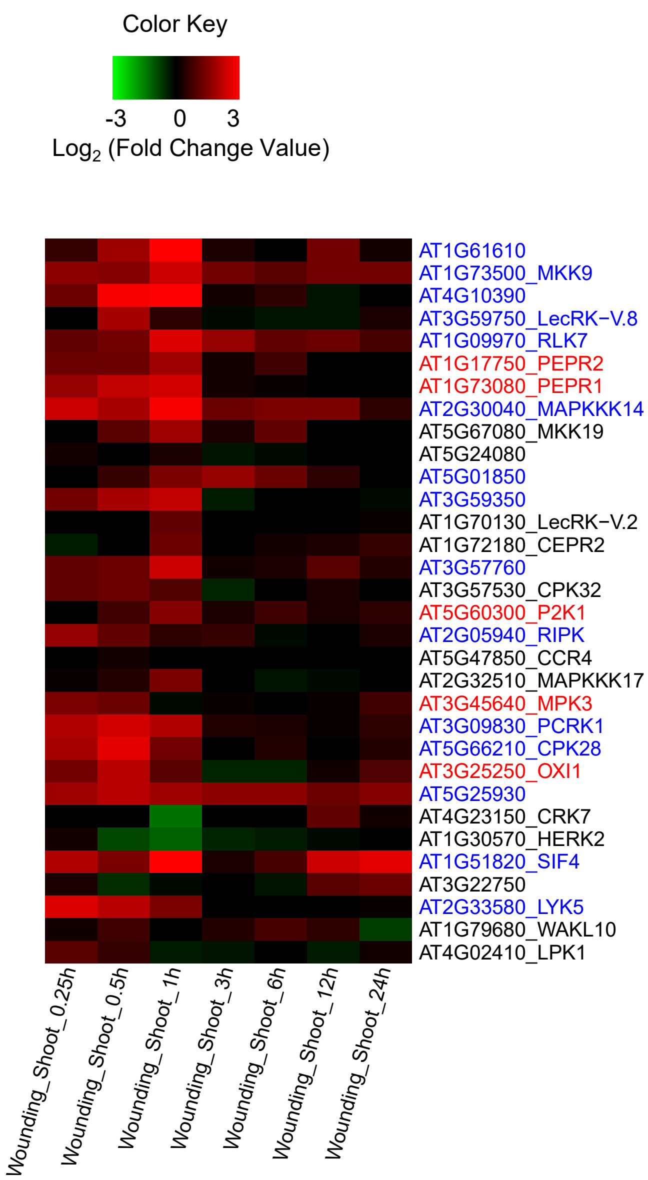

### Fig. S4

Figure S4

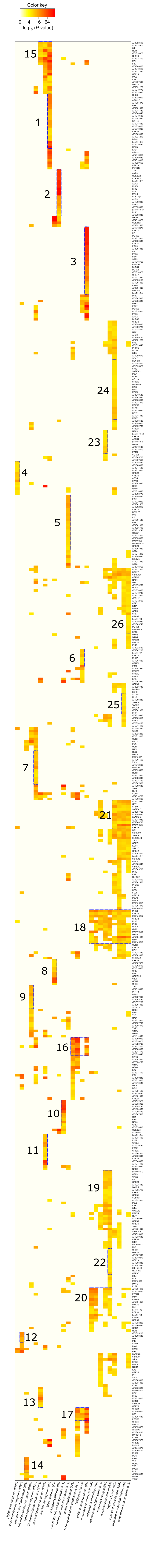
